# The ultrastructural nature of human oocytes’ cytoplasmatic abnormalities and the role of cytoskeleton dysfunction

**DOI:** 10.1101/2023.04.20.537668

**Authors:** Martina Tatíčková, Zuzana Trebichalská, Drahomíra Kyjovská, Pavel Otevřel, Soňa Kloudová, Zuzana Holubcová

**Affiliations:** Department of Histology and Embryology, Faculty of Medicine, Masaryk University, Brno, Czech Republic; Reprofit International, Clinic of Reproductive Medicine, Brno, Czech Republic

**Keywords:** human oocyte, dysmorphism, oocyte abnormalities, refractile bodies, electron microscopy, egg quality, cytoskeleton, actin

## Abstract

**Background:** Egg quality is a limiting factor of female fertility and assisted reproductive technology (ART) success. Oocytes recovered from hyperstimulated ovaries often display morphological anomalies suspected to compromise their fertilization and developmental potential. Knowledge of (ab)normal oocyte’s intracellular organization is vital to establish reliable criteria for morphological evaluation of oocytes intended for in vitro fertilization (IVF).

**Methods:** Transmission electron microscopy (TEM) was used to investigate the fine morphology of 22 dysmorphic IVF eggs exhibiting different types of cytoplasmic irregularities, namely (1) refractile bodies, (2) centrally-located cytoplasmic granularity (CLCG), (3) smooth endoplasmic reticulum (SER) disc, and (4) vacuoles. The cytoskeleton targeting compounds were employed to address the causative mechanism behind the anomalous cytoplasmic architecture observed in abnormal egg samples. A total of 133 immature oocytes were exposed to chemical inhibitors/control conditions, and their morphology was examined by fluorescent and electron microscopy.

**Results:** TEM exposed the structural basis of the common oocyte aberrations and revealed that the underlying cause of two of the studied morphotypes was excessive organelle clustering. Inhibition experiments showed that disruption of actin, not microtubules, allows inordinate aggregation of subcellular structures resembling the ultrastructural pattern seen in morphologically abnormal eggs retrieved in IVF cycles. These results imply that actin serves as a regulator of organelle distribution during human oocyte maturation.

**Conclusions:** The ultrastructural analogy between dysmorphic eggs and oocytes, in which actin network integrity was perturbed, suggests that malfunction of the actin cytoskeleton might be implicated in generating common cytoplasmic aberrations. Knowledge of human oocytes’ inner workings and the origin of morphological abnormalities is a step forward to more objective egg quality assessment in clinical practice.

## BACKGROUND

Eggs are commonly reffered to as good or bad according to their chromosomal content, while the quality-determination role of its enormously large cytoplasm is less recognized. As the oocyte grows, the cytoplasm stockpiles cellular material and molecular factors needed to support embryo metabolism after fertilization. The final stage of egg development, oocyte maturation, is marked by structural and biochemical modifications, rendering the oocyte capable of completing meiotic segregation, fertilization, and early embryogenesis (1–3). Human eggs are well-known to vary in their fertilization and developmental competence. The technical and ethical issues related to the provision of human oocytes for research are limiting the study of the fundamental biology of these unique cells and exploration of factors affecting their quality.

In ART practice, female gametes are harvested from preovulatory follicles, and morphology of denuded oocytes intended for intracytoplasmic sperm injection (ICSI) or vitrification is evaluated by conventional stereomicroscopic examination. Preovulatory oocytes retrieved in controlled ovarian stimulation (COS) cycles tend to differ in their morphological appearance, maturation grade, and developmental capacity. The good-quality mature egg is characterized by a spherical shape, a single, normal-sized polar body (PB) and perivitelline space (PVS), a uniform zona pellucida (ZP), and a pale cytoplasm with homogenous texture and smooth appearance (4). However, the oocytes derived from IVF patients often deviate from this ideal picture. Hormonally-primed follicles might occasionally contain giant or misshaped oocytes. Remarkably big or fragmented PB, atypical ZP, and large or small PVS are collectively termed extracellular defects. The most common cytoplasmic irregularities include refractile bodies, increased granularity of ooplasm, and vacuoles. On the other hand, a smooth disc-shape structure disrupting cytoplasmic texture in phase contrast is encountered only occasionally. An overview of dysmorphic phenotypes is available in the literature, but their biological significance is unclear (5–7).

Multiple studies aimed to evaluate the relationship between oocyte morphology and IVF outcome. However, the published evidence is controversial. Some authors reported that abnormal oocytes have a lower chance of producing transferable embryos and healthy pregnancies (8–12), but others found no correlation between oocyte morphology and its developmental capacity (13–16). The conflicting data might be explained by inconsistent dysmorphic patternś classification and different study endpoints. Some investigators focused on a single morphotype only, whereas others considered multiple abnormal patterns. The interpretation of results is further complicated by the fact that multiple cellular aberrations may coincide in one cell; thus, their individual impact on oocyte quality cannot be readily dissected (17). Meta-analysis, which drew together results from 14 studies, concluded that the probability of an oocyte becoming fertilized is significantly reduced by the presence of a large PB, large PVS, refractile bodies, and vacuoles (18). An international consent meeting of ART experts attempted to set standards for egg and embryo features assessment in clinical practice (4). Nevertheless, there is still considerable ambiguity regarding the definition and predictive value of distinct morphotypes. Without clearly defined classification criteria and widely accepted guidelines, no deselection of observed morphological abnormalities is routinely applied in clinical laboratories, and all collected metaphase MII (MII) oocytes are commonly used for ICSI or cryopreservation.

Unravelling the structural bases of morphological anomalies is necessary to determine their impact on oocyte metabolism and developmental competence. Nevertheless, the documentation of dysmorphic egg ultrastructure is scarce and scattered in literature. In 1990, Van Blerkom was the first who employed TEM to analyze human oocytes exhibiting aberrant cytoplasmic features (19). His micrographs revealed that large rounded cytoplasmic inclusion with a smooth and flat appearance is not a plain vacuole, but an enormous aggregate of smooth endoplasmic reticulum (SER), thereby referred to as a SER disc. The structural character of this anomaly was confirmed by later studies (11, 20), which also reported that SER-positive oocytes had reduced developmental capacity. A combination of TEM and spectral imaging showed that refractile bodies, visible as dark specks in phase contrast, are heterogenous clusters of fibrous material, granular vesicles, and electron-dense lipid substances exerting characteristic autofluorescence (19, 21). Interestingly, granule-fibrillar inclusions were also found in the interior of atypical granular vacuoles (6). Collectively, previously published reports cast light on the ultrastructural nature of the common oocyte abnormalities. However, they did not cover a full spectrum of dysmorphic phenotypes, and the number of TEM-imaged samples is small.

Little is known about the etiology of human oocyte morphological anomalies. Lacking robust scientific evidence, our views are shaped by anecdotal evidence and incidental findings combined with theoretical presumptions. The diminished quality of female gametes was linked to an atypical hormonal profile in IVF patients (20, 22). Therefore, oocyte dysmorphisms are believed to arise as a suboptimal response to the stimulation regimen. Van Blerkom was fortunate to capture the rapid formation of the endocytic vacuole in fertilized human ova and proposed that oocyte vacuolization is caused by an instability of the cellular cortex (19). Observing the growth of the SER disc during prolonged cultivation of uninseminated eggs Otsuki and colleagues assumed that it is a sign of aging-related cellular deterioration (20). The popular review postulated that cytoplasmic granularity is attributed to excessive clustering of oocyte organelles (5). Nevertheless, ultrastructural data supporting this notion are missing.

Oocyte dysmorphisms appear to derive from alteration in the morphologically normal progression of cytoplasmic development during preovulatory oogenesis. However, no studies tackled to address molecular mechanisms underlying the emergence of different cytoplasmic irregularities in developing human oocytes. We have previously mapped out ultrastructural changes during human oocyte maturation, and our 2D and 3D image data documented major reorganization of ooplasm in preparation for fertilization (23, 24). Correct positioning and active movement of organelles within the cytoplasm rely on their interaction with cytoskeleton components. Thus, actin/microtubule deficiency to orchestrate organelle relocation during meiotic maturation may underlie the emergence of cytoplasmic abnormalities in IVF oocytes. However, the role of the cytoskeleton in the human oocyte cytoplasmic maturation and the development of dysmorphic features was not experimentally verified.

In this study, we harnessed our experience with electron microscopy of normal human oocytes (23, 24) and examined the subcellular morphology of dysmorphic female gametes rejected for ICSI. To elucidate cellular mechanisms implicated in the evolution of cellular aberrations, we experimentally perturbed cytoskeleton function in oocytes maturing in vitro and inspected their inner morphology.

## MATERIAL AND METHODS

### Source of oocytes

A total of 150 human oocytes from 102 young and healthy egg donors and 5 oocytes from 3 IVF patients were analyzed in this study (Supplementary Table). Women enrolled in the clinical egg donation program underwent screening for hormonal and genetic factors that could negatively affect reproduction. Ovarian stimulation and egg retrieval were performed according to the established clinical protocol described in detail previously (23). The cumulus cells were enzymolyzed in hyaluronidase (90101, Irvine Scientific), and the meiotic status of each oocyte was determined under stereomicroscope based on the presence/absence of the germinal vesicle (GV)/the first polar body (PB). Surplus oocytes unsuitable for ICSI were used for research only if the donor’s written informed consent was obtained. The study was undertaken under ethical approvals issued by the Ethics Committees of the collaborating academic institution and clinical unit.

### Oocyte cultivation and cytoskeleton inhibition

Dysmorphic eggs rejected for IVF were handed over for research after denudation and fixed without further delay. The spare immature oocytes were cultured until noon the next day in the maturation medium (G-MOPS, Vitrolife) at 37 °C and 5 % oxygen. Perturbation of cytoskeleton during in vitro maturation was achieved by overnight exposure of morphologically normal GV oocytes to drugs known to interfere with the function of actin network (1 µM cytochalasin D, C8273, Sigma Aldrich; 5 µM brefeldin A, B7651, Sigma Aldrich) and microtubules (1 µM paclitaxel, T7402, Sigma Aldrich; 33 µM nocodazole, M1404, Sigma Aldrich, 3 different commercial lots tested). The effect of nocodazole during overnight/short-term culture was enhanced by 10 µM verapamil (V4629, Sigma Aldrich). To verify microtubule depolymerization, the presence of bipolar meiotic spindle was first confirmed by polarized light microscopy (PLM) as previously described (25). Then, MII oocytes were subjected to a 1-hour drug treatment followed by fixation. Control cells were exposed to the corresponding concentrations of dimethyl sulfoxide (DMSO). All inhibition experiments were carried out without oil overlay to avoid disproportional drug distribution into oil phase. Stable humidity was ensured by filling an inter-well space with water.

### Electron microscopy

For TEM inspection, oocytes were fixed in 3% glutaraldehyde (G5882, Sigma Aldrich) in 0.1M cacodylate buffer (C0250, Sigma Aldrich) (pH 7,2 – 7,8) at room temperature overnight. Next, oocytes were post-fixed with 1% osmium tetroxide (O5500, Sigma Aldrich) in 0.1 M sodium cacodylate buffer supplemented with 1.5% potassium ferrocyanide (1049821000, Sigma Aldrich) for 1 hour. Individual samples were mounted into agarose, dehydrated and embedded in epoxy resin, as described before (23) . Ultrathin sections placed on grids were stained with 1% aqueous uranyl acetate (Agar Scientific, Stansted, UK) and 3% lead citrate (Sigma Aldrich, St. Louis, USA), and observed at the FEI Morgagni 268D microscope.

### Fluorescence microscopy

To assess meiotic spindle and chromosome configuration following drug treatment, individual oocytes were fixed for 60 min at 37°C in 100 mM HEPES (pH 7) titrated with KOH, 50 mM EGTA (pH 7) titrated with KOH, 10 mM MgSO_4_, 2% formaldehyde (MeOH free) and 0.2% Triton X-100, based on previously published protocol (26). After fixation, oocytes were rinsed in phosphate-buffered saline supplemented with 0.1% Triton X-100 (PBT) and kept in PBT overnight at 4°C. For microtubule staining, oocytes were exposed to an anti-α-tubulin antibody (rat monoclonal MCA78G, Bio-Rad) overnight at 4°C and Alexa-Fluor-633-labelled goat anti-rat antibody (A-21094, Thermo Fisher Scientific) for 2 hours in the dark and at room temperature. Chromatin was stained by DAPI (D1306, Thermo Fisher Scientific) and actin filaments by Alexa-Fluor-488 labeled phalloidin (A12379, Thermo Fisher Scientific) for 1 hour in the dark and at room temperature. Incubation with primary/secondary antibody and fluorescent dyes were carried out in PBT supplemented with 3% bovine serum albumin. For mitochondria visualization, live cells were treated with the fluorescent probe MitoTracker Orange CM-H2TMRos (M7511, Thermo Fisher Scientific) for 1 hour in the dark and at 37 °C before fixation. Samples were imaged using Zeiss LSM 800 confocal fluorescence microscope.

## RESULTS

### Ultrastructural analysis of dysmorphic oocytes

The fine morphology of 22 dysmorphic oocytes, which exhibited prominent cytoplasmic abnormalities during stereo microscopy examination, was inspected by TEM. The samples were derived from 3 IVF patients (30,33 and 42 years old) and 14 healthy egg donors (mean age 28.86 ± 6.20 years). The refractile bodies were the most common cytoplasmic inclusions in our samples (18 out of 22 oocytes). Vacuoles were observed in 11 samples, and cytoplasmic granularity in 6 samples. Although only 1 oocyte exhibited SER disc under low magnification, TEM analysis revealed enlarged SER clusters in another 3 oocytes. In 9 oocytes only one type of dysmorphism was present, while in 13 oocytes two or three different defects coincided (Supplementary table).

### Refractile bodies

Refractile bodies were visible in the light microscope as prominent various-sized and typically dark specks disrupting cytoplasm homogeneity. Small refractile bodies (2–5 μm) occurred either isolated or alongside other irregularities, and were typically localized in the central part of the cytoplasm (Figure 1A). Interestingly, 3 examined eggs featured extremely large refractile bodies which exceeded 10 μm in diameter and appeared lighter under stereomicroscope (Figure 1 D). When viewed under the electron microscope, the affected oocytes exhibited pleomorphic bodies depositing electron-dense granular material, amorphous substances, vesicles, lipid droplets, and membrane remnants (Figure 1 B, C, E, F-N). These clumps of heterogeneous cellular debris were typically delimited by a discontinuous membrane and surrounded by cytoplasm populated by oocyte mitochondria with atypical arch-like or transversal cristae, tubular and vesicular type of SER, and small vesicles (Figure 1 B, C, E-N), as previously described in human oocytes deemed morphologically normal by established clinical criteria (23). Notably, even the presence of oversized refractile bodies did not alter the homogeneous distribution of organelles characteristic for MII oocytes and maturation-induced formation of “necklace” complexes composed of multiple mitochondria attached to individual SER sacs (23) (Figure 1 B, C, L, N).

**Figure 1:**
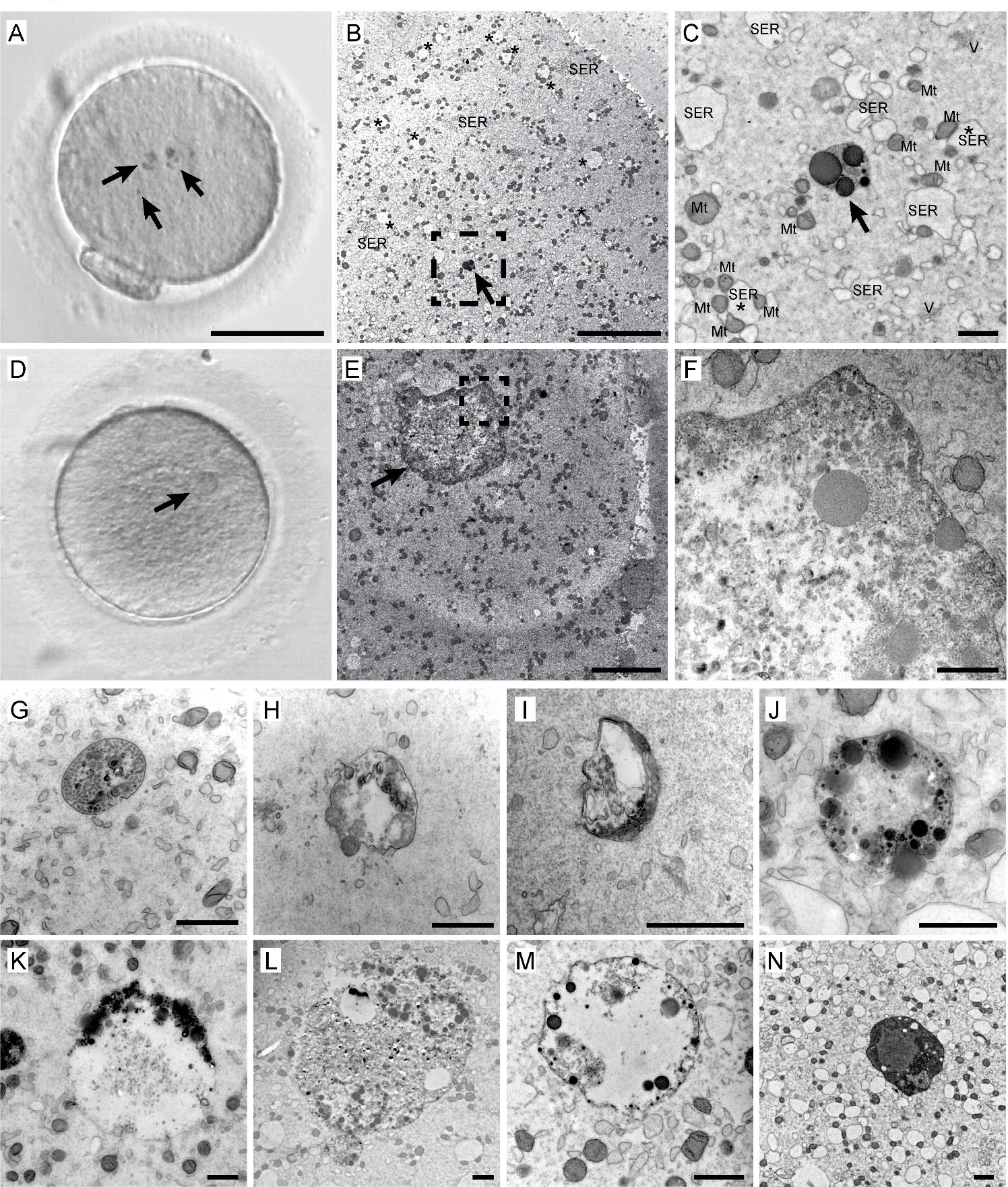
Examples of refractile bodies observed in dysmorphic human eggs. Live cell appearance in transmitted light before fixation (A, D) and TEM micrographs of the two representative MII oocytes showing different numerous small (A) or one big inclusion (D). Overviews (B, E) and details (C, dashed rectangle in B; F, dashed rectangle in E) of the corresponding cellś ultrastructure. Refractile bodies are indicated by arrows (A-E); typical organelles are labelled: Mt-mitochondria, SER – smooth endoplasmic reticulum, V – vesicles; asterisks indicate Mt-SER („necklace“) complexes characteristic of mature human eggs (B, C). Overview of ultrastructural variability of refractile bodies found in multiple sample cells (G-N). Membrane-bound bodies exhibit variable content, which could include granular material (G, H, J-N), membrane remnants (G-I), and lipid droplets (J-N). Scale bar, 50 μm (A, D), 10 μm (B, E), 1 μm (C, F, G-N).

### Cytoplasmic granularity

Centrally-located cytoplasmic granularity (CLCG) was identified during the routine stereomicroscopic examination as a large (> 30 μm in diameter) crater-like region residing in the celĺs center (Figure 2 A). Ultrastructural analysis of 6 CLCG-exhibiting eggs showed that the uneven texture of their cytoplasm was caused by anomalous organelle distribution. The large granular area in the cell center featured an accumulation of enlarged SER sacs attended by closely associated mitochondria, vesicles, and lysosomes. In contrast, peripheral cytoplasm was harbored only individual mitochondria and tiny SER vesicles (Figure 2 B, C). Most SER cisternae within the central organelle conglomerate were markedly dilated compared to peripheral ones, and some reached the size of small vacuoles. The SEŔs membrane appeared undulated and confined clear fluid, occasionally containing small granules with an electron-dense substrate. Numerous small refractile bodies appeared trapped in dense organelle assemblages (Figure 2 B, C).

**Figure 2:**
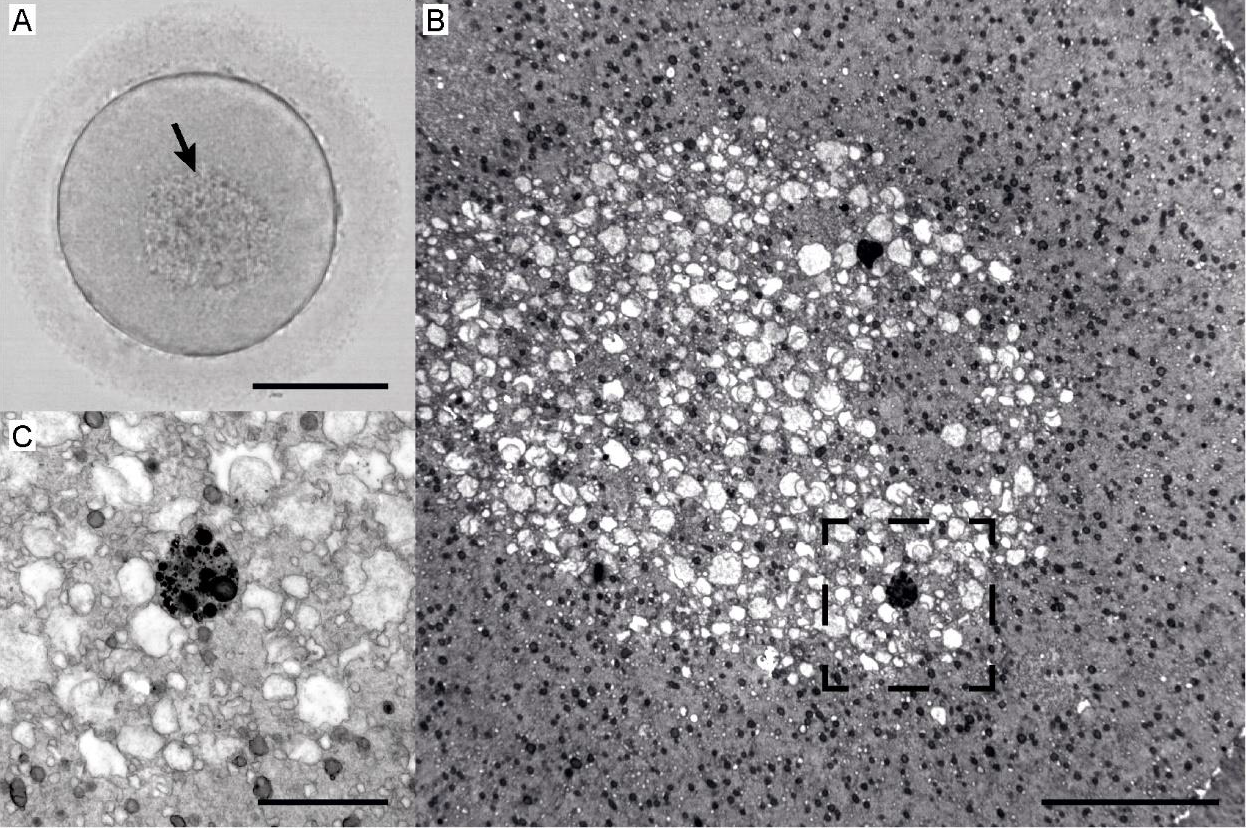
Example of a human egg exhibiting prominent centrally-located cytoplasmic granularity (CLCG) Live cell appearance in transmitted light before fixation (A, CLCG indicated by arrow) and TEM micrographs (B, C) of its ultrastructure. Overview of ooplasm with excessive organelle clustering in the cell center (B) and magnified detail of organelle cluster periphery (dashed area) presenting a refractile body surrounded by dilated SER cisternae, mitochondria, and small vesicles (C). Scale bar, 50 μm (A), 10 μm (B), 2 μm (C).

### SER disc and large SER aggregates

The rare cytoplasmic abnormality, known as SER disc, was recognized in 1 egg in our sample cohort. This cytoplasmic aberration appeared in phase contrast as a flat, smooth plaque with an indistinct outline and diameter exceeding 30 μm (Figure 3 A). Ultrastructural analysis showed that the vacuole-like structure was a giant assemblage of elongated tubular-type SER. The fine meshwork of curvilinear tubules was free of other organelles. Due to absence of the membrane, the SER mass periphery was in direct contact with the surrounding cytoplasm (Figure 3 B, C, Supplementary Figure 1B). Adjacent to the SER disc was a small vacuole containing electron-dense granules and lipid droplets (Figure 3B). Surrounding cytoplasm was found to be occupied by mitochondria-SER complexes indicating egǵs maturity (Figure 3B, Supplementary Figure 1). Along with a massive SER disc and small “necklace” complexes, we also observed mid-sized (∼5-7 μm) clusters of densely arrayed tubular SER decorated by only a few mitochondria (Supplementary Figure 1). Small refractile bodies were scattered in the cytoplasm (Supplementary Figure 1 C). The meiotic spindle poles appeared to be loosened and chromosomes misaligned, indicating that the developmental potential of this particular oocyte was impaired (Supplementary Figure 1C). Interestingly, mid-sized SER clusters, undiscernible under low magnification, were found in another 3 oocytes analyzed for different dysmorphisms (Figure 4 B, Supplementary Figure 2, Supplementary Figure 3).

**Figure 3:**
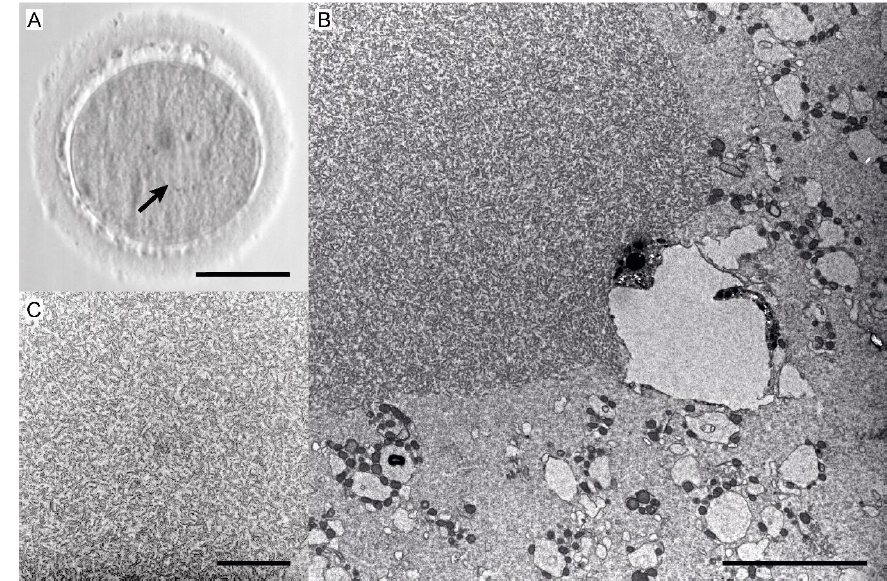
Human egg exhibiting SER disc. Live cell appearance of affected MII oocyte in transmitted light before fixation (A, arrow indicates smooth rounded area corresponding to SER disc) and in TEM micrographs (B, C) of its ultrastructure. Overview of SER disc with adjacent granular vacuole and surrounding cytoplasm (B) and detail of SER disc interior. (C). Scale bar, 50 μm (A), 5 μm (B), 2 μm (C).

**Figure 4:**
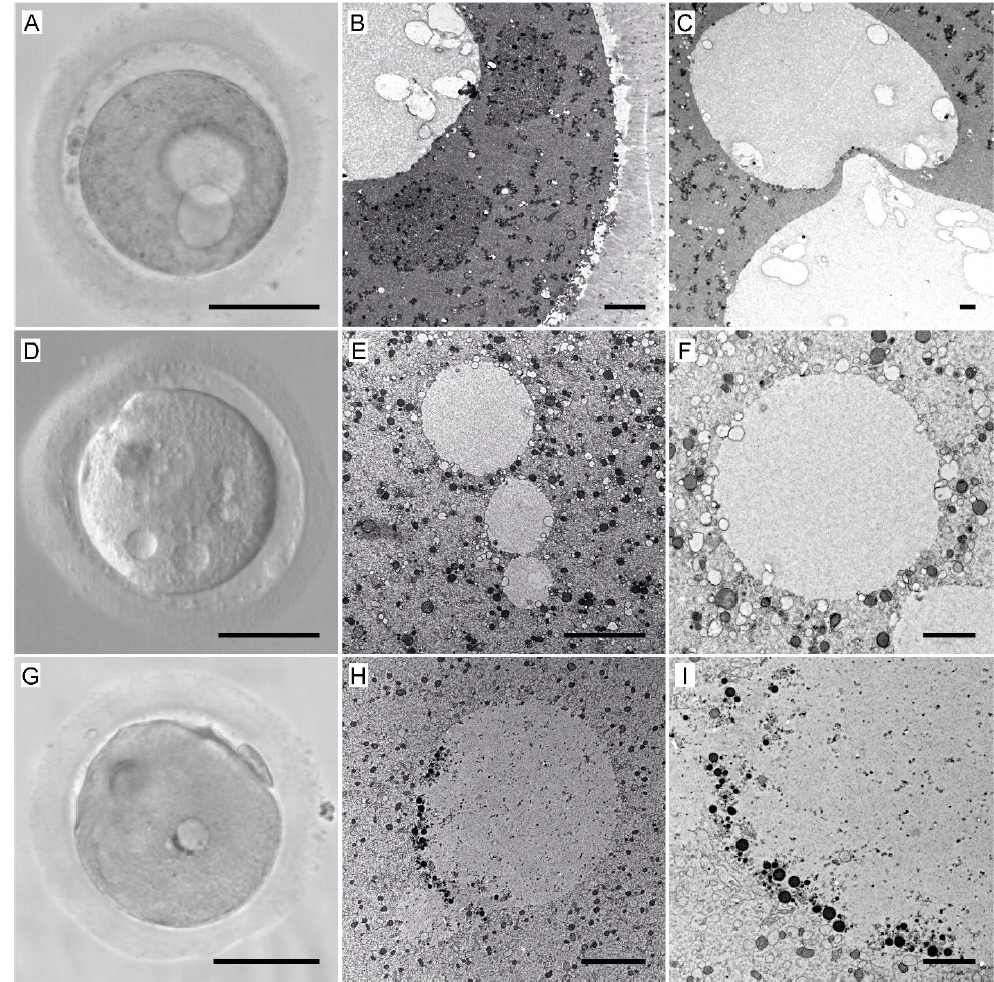
Examples of oocytes exhibiting vacuoles. First oocyte with two large vacuoles (A-C), second one with multiple small vacuoles (D-F), and third one a granular vacuole (G-I): live cell appearance in transmitted light before fixation (left column) and TEM micrographs – overview (middle column) and detail (right column) of the corresponding cells. Scale bar, 50 μm (A, D, G), 5 μm (B, E, H), 2 μm (C, F, I).

### Vacuoles

The severe vacuolization led to the ICSI rejection of 11 MII oocytes. The vacuoles differed in size, number, and appearance (Figure 4). In contrast to SER disc, all vacuoles were clearly visible in transmitted light and had well-defined boundaries (Figure 4 A, D, G). Inspection of oocyte morphology under high magnification confirmed that these round reflective cavities were enclosed by a membrane and filled with fluid with a translucent or slightly floccular appearance (Figure 4 B, E, H). A few TEM images captured adjacent vacuoles with their membranes in intimate contact, suggesting that these dynamic organelles might be prone to merging (Figure 4 C, E, F, H, I). One oocytés vacuole was asymmetrically lined with fat droplets, and fine granules were present in its interior (Figure 4 G, H, I). Moreover, the oocyte featuring two enormous vacuoles harbored horseshoe- or ring-shaped mitochondria dispersed in the ooplasm, numerous little vacuoles, and sizable patches of amassed tubular SER surrounded by multiple mitochondria (Figure 4 B, Supplementary Figure 3 A, B). While SER disc and mid-sized aggregates detected in other oocyte samples were exclusively composed of tubular SER elements, here, the dense SER assemblage encompassed electron-dense granules, mitochondria, and small vacuoles (Figure 4 B, Supplementary Figure 3 A). Instead of an MII spindle with individualized chromosomes, we located condensed genetic material, which collapsed into an amorphous lump (Supplementary Figure 3 C). Together, observed morphological features denoted that the affected oocyte was pathological.

### Perturbation of cytoskeleton

To test the hypothesis that some cytoplasmic irregularities may arise from cytoskeleton dysfunction, we targeted actin and microtubule function in oocytes maturing in vitro. A total of 88 GV oocytes from 56 hormonally stimulated women enrolled in the clinical egg donation program (Supplementary table) were incubated overnight with small-molecule cytoskeletal inhibitors and examined by light, immunofluorescence, and electron microscopy (Figure 5 A). Specifically, were evaluated the maturation efficiency, global distribution of organelles, the integrity of the oolemma, and the chromosome-spindle configuration. The observed phenotypes were compared with control samples cultured simultaneously in maturation medium supplemented with DMSO (45 oocytes from 32 donors). Besides, our archive of TEM micrographs of morphologically normal in vitro-matured oocytes (23) was used as a reference.

**Figure 5:**
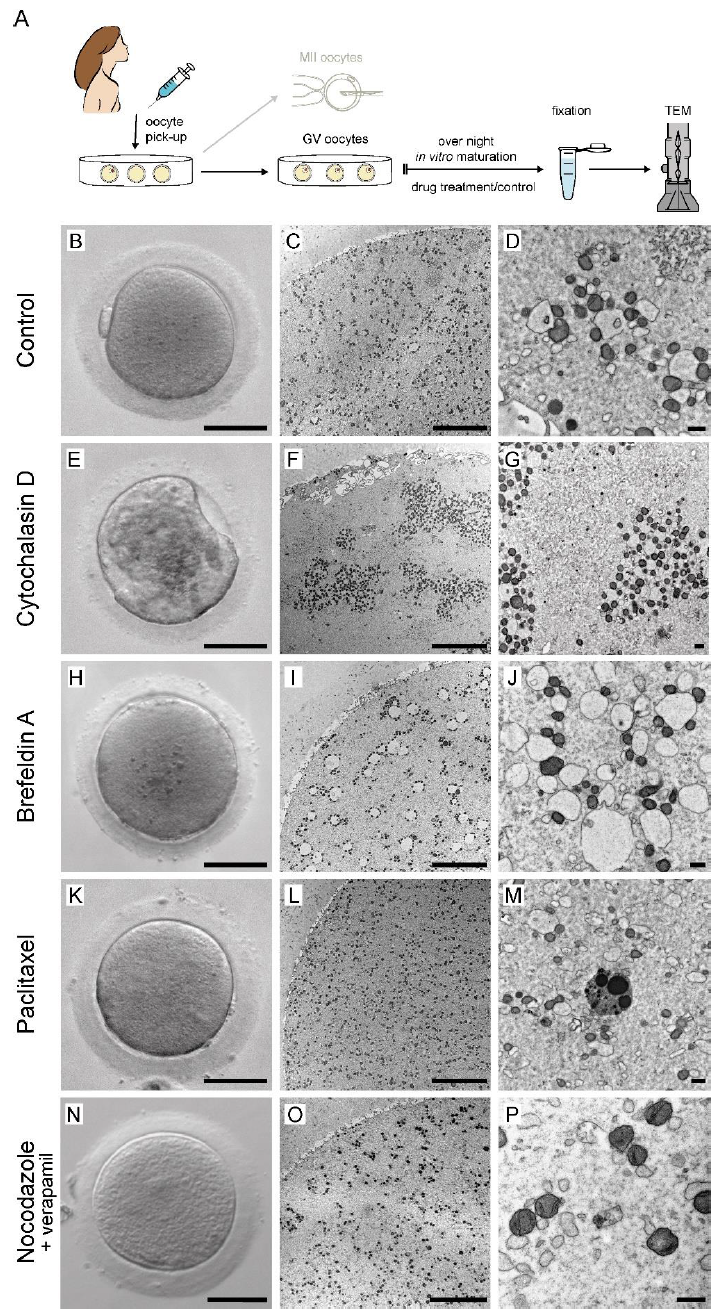
Perturbation of cytoskeleton. Schematic depiction of experimental set-up (A). GV oocytes retrieved in donor IVF cycles were cultured overnight in presence of cytoskeleton-affecting drugs or corresponding concentration of DMSO. Live cell appearance in transmitted light (left column) and TEM micrographs of the same cell showing overviews (middle column) and details (right column) of control (B-D) and drug treated-oocytes (E-P). Cytochalasin D (E-G) and brefeldin A (H-J) were used to inhibit the actin network. Paclitaxel (K-M) and nocodazole enhanced with verapamil (N-P) were used to inhibit microtubules. Granular cytoplasm and excessive organelle clustering is visible in oocytes in which actin polymerization was blocked (E-G). Scale bar, 50 μm (B, E, H, K, N), 10 μm (C, F, I, L, O), 1 μm (D, G, J, M, P).

An inhibitor of actin polymerization, cytochalasin D (CytD), was applied to disrupt actin network organization. Unlike control cells, which showed normal maturation rate, overall morphology, chromosome configuration, and ultrastructural pattern characteristic for human eggs (Supplementary Figure 4, Figure 5 B-D), all 14 oocytes matured in the presence of CytD failed to extrude PB after overnight culture and were grossly dysmorphic (Supplementary Figure 4 A, Figure 5 E). Fluorescence microscopy confirmed that both the cell cortex and non-cortical network of actin filament were impaired and non-homogenous distribution of organelles explained irregular cytoplasmic texture. The meiotic spindle was aberrant and misplaced (Supplementary Figure 4 B). When viewed in electron microscopy, CytD-treated cells featured massive mosaic clusters of mitochondria, SER cisternae, and small vesicles. In addition, TEM revealed the presence of massive aggregates of loosely arrayed tubular SER (Figure 5 F, G). The plasma membrane of CytD-treated oocytes appeared ruffled and devoid of microvilli. Sectioning through the cortical area showed that cortical granules grouped beneath the oolemma but were sparse in comparison to controls (Figure 5 F).

In mouse oocytes, Brefeldin A (BFA) was found to inhibit actin network dynamics without affecting its structural integrity (27). Thus, we decided to expose immature human oocytes to this modulator of actin dynamics to determine whether structural or functional properties of the actin network are required for homogenous organelle distribution. In contrast to CytD-treated oocytes, oocytes cultured in presence of BFA retained the moderate capacity to complete maturation (6 out of 18 oocytes) and showed no prominent morphological anomalies (Supplementary Figure 4 A, Figure 5 H). The meiotic spindle was located at the cortex but was apolar and carried misaligned chromosomes (Supplementary Figure 4 B). Although the organelle distribution appeared normal in fluorescent images, the ultrastructural analysis revealed that SER cisternae within “necklace” SER-mitochondria complexes were markedly swollen (Figure 5 I, J). Dilatation of SER was particularly prominent in prophase-arrested GV oocytes, and phenotype receded as maturation progressed (Supplementary Figure 5). Unlike in the CytD group, the plasma membrane of BFA-treated oocytes showed no obvious pathology and was covered with microvilli (Figure 5 I). Compared to control conditions, an increased number of tiny refractile bodies was observed in the cytoplasm. A striking difference between the CytD and BFA phenotypes suggests that structural integrity of actin filaments rather than vesicle-driven network dynamics is required for homogenous organelle distribution during cytoplasmic maturation.

Next, we sought to determine the role of microtubules in the rearrangement of ooplasm occurring during human oocyte maturation (23). Two anti-mitotic drugs were used to evaluate if the inhibition of microtubules can induce excessive organelle clustering. As expected, the microtubule-stabilizing drug paclitaxel (PX) application prevented chromosome segregation in 9 out of 11 meiotically competent oocytes (Supplementary Figure 4 A). In fluorescently stained samples, the spindle apparatus was disorganized and detached from the plasma membrane (Supplementary Figure 4 B). Nevertheless, the overall oocytés shape and cytoplasmic texture appeared normal (Figure 5 K). Neither fluorescent nor electron microscopy uncovered deviation from a normal distribution of organelles (Supplementary Figure 4 B, Figure 5 L). Fine morphology of the oolemma and adjacent cortical area showed no irregularities. Tiny refractile bodies, non-discernible under the stereomicroscope, were observed in TEM images of all PX-treated oocytes (Figure 5 M).

To our surprise, the application of nocodazole (NC), known as a potent inhibitor of microtubule polymerization, did not prevent meiotic spindle assembly and PB extrusion in our experiments. Out of 22 GV oocytes exposed to NC during overnight culture, 21 cells assembled bipolar spindle, and 15 completed their maturation by midday the next day (Supplementary Figure 4A, Supplementary Figure 6 A). Thus, we decided to probe the anti-tubulin effect by short-term treatment of 5 MII oocytes exhibiting a birefringent spindle. However, the immunofluorescence imaging showed that the meiotic spindle remained insensitive to 1-hour NC exposure (Supplementary Figure 6 B). The rapid spindle disturbance was observed when the calcium channel blocker, verapamil, was added to the maturation medium (Supplementary Figure 6 A, B). During overnight culture, 10 out of 11 GV oocytes resumed meiosis, but only 1 oocyte extruded a PB. These experiments demonstrated that combined drug treatment did not impede meiotic resumption but efficiently prevented chromosome segregation (Supplementary Figure 4 A). All oocytes incubated in the presence of NC and verapamil showed normal appearance in phase contrast, and no alteration in organelle distribution and morphology was found at the ultrastructural level. Also, membrane and cortical region architecture did not differ from the control cells (Figure 5 N-P).

## DISCUSSION

Electron microscopy proved to be a powerful tool enabling investigation of the fine morphology of human oocytes, which are available for research only in small numbers. Detailed inspection of subcellular organization provides information about the nature of morphological aberrations indicated by routine oocyte examination. Moreover, high-magnification scrutiny can reveal features that are undiscernible at the light-microscopy level. In this study, we performed TEM analysis of IVF oocytes with distinct cytoplasmic abnormalities and evaluated the degree of their deviation from normal oocytes’ ultrastructural pattern (Figure 6). Furthermore, we demonstrated that disruption of the actin network in maturing oocytes leads to a similar misarrangement of subcellular structures as seen in dysmorphic eggs.

**Figure 6:**
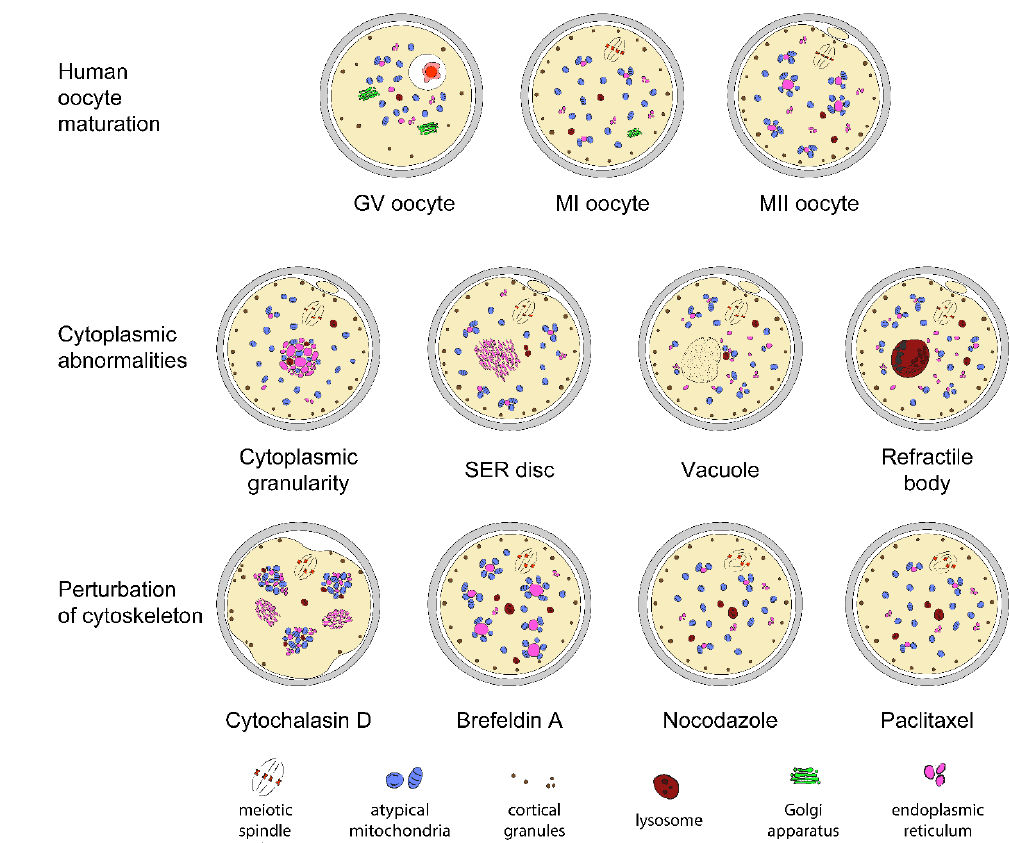
The schematic representation of the subcellular morphological organization of normal oocytes, dysmorphic oocytes, and oocytes with the perturbated cytoskeleton.

The morphological irregularities are common in human oocytes retrieved from hormonally stimulated ovaries. Here, the supraphysiological concentration of gonadotropins enhances the recruitment of subordinate follicles and rescues them from atresia. According to a study involving 516 ICSI cycles, over 90% of patients had at least one abnormal oocyte (10), and the overall incidence of oocyte dysmorphisms across studies is reported as high as 60-80% (12-14, 16, 28). Some anomalies may be attributed to intrinsic factors such as genetic background and patient’s age, while others seem to be related to the suboptimal intrafollicular environment (e.g., oxidative stress, hypoxia, proapoptotic factors, inflammation, and hyperglycemia). Recently breast cancer was identified as a risk factor for the presence of dysmorphic oocytes (29). This evidence suggests that alteration of hormonal receptors and/or signaling pathways might impair follicle-oocyte dialogue and compromise the quality of developing female gametes. Previous ultrastructural studies analyzed only dysmorphic oocytes derived from infertile patients (6, 11, 19, 20, 30). We also document the most common cytoplasmic abnormalities in oocytes from young women with no reproductive issues. In IVF patients, cellular perturbations often affect all oocytes collected and show a high rate of recidivism (31, 32). In egg donation program, severe aberrations are seen only as a minority of the cohort, illustrating a quality hierarchy of hormone-responsive follicles. Intra-ovarian regulation ensures mono-ovulation of the best available egg in our species. But administration of exogenous hormones overrides this quality control mechanism unmasking phenotypic variability of inferior female gametes that are destined to apoptosis under physiological conditions. All dysmorphic oocyte samples in this study were fixed shortly after denudation. Therefore, adverse effects of in vitro ageing can be excluded.

Refractile bodies were the most frequently observed type of cytoplasmic abnormalities in our samples. The detailed analysis showed that these inclusions are heterogeneous clumps of degraded cellular material, which is in line with published TEM reports (19, 30). Our experience that tiny electron-dense refractile bodies can be detected even in human oocytes with normal appearance (23) implies that this irregularity represents phenotypical variation rather than detrimental cytopathology. The ultrastructure of refractile bodies resembles tertiary lysosomes storing cellular waste. These catabolic organelles are abundant in post-mitotic cells in which the lysosomal degradation pathway ensures homeostasis and lifespan control (33). Similarly, membrane-bound refractile bodies may deposit undegradable substances such as products of proteolytic activity, lipid peroxidation, phagocytosis, and autophagy to avoid the buildup of unwanted and damaged cellular material in the cytoplasm of long-lived human oocytes. This notion is supported by the evidence that large refractile bodies are loaded with oxidized lipochrome lipofuscin (30). This insoluble pigment accumulates over time in neurons, cardiac and skeletal muscle, retinal pigment epithelium and hepatic cells, and is considered as the hallmark of aging (34). The hypothesis that refractile bodies incorporate biological “garbage” and/or sequester xenobiotics accumulated in terminally differentiated cells throughout a woman’s life is endorsed by clinical experience that the incidence of dark inclusions is age-dependent (unpublished data). Whether the abnormal protein or lipid metabolism, reduced capacity of intralysosomal degradation, oxidative stress, or intracellular deposition of insoluble toxic materials are involved in the generation of refractile bodies remains to be explored.

SER disc is regarded as the most severe cytoplasmic abnormality of human oocytes. Clinical data indicate that embryos derived from affected oocytes have a higher risk of suboptimal IVF outcomes, including a reduced chance of pregnancy, obstetric complications, and congenital malformations (11, 20, 31, 35). Therefore, professional authorities strongly recommended that oocytes displaying a SER disc should not be used for ICSI (4). Nevertheless, there are reports that SER disc-positive oocytes can develop normally and give rise to healthy newborns (36, 37). The SER disc could be mistaken for a vacuole during the routine morphological assessment. But at the ultrastructural level, a massive assemblage of SER is easily distinguishable from the membrane-bound fluid-filled vacuoles, as demonstrated by us and others (19, 20). This study involved one oocyte exhibiting a single pronuclear-sized SER disc. Unlike in earlier studies, here, the SER disc periphery was not lined with mitochondria which could explain the structurés poor visibility during routine oocyte inspection. Notably, the ultrastructural analysis showed that sizeable SER aggregates also resided in 3 more oocyte samples which were scrutinized due to different types of dysmorphism (Supplementary Table). We have previously monitored structural changes during cytoplasmic maturation and observed the progressive association of mitochondria with SER and the formation of heterotypic SER-mitochondria complexes in oocytes with normal appearance. In addition to characteristic “necklace” complexes, composed of a SER sac surrounded by a corona of mitochondria, small aggregates of tubular SER (≤ 3 μm) decorated by a few mitochondria were detected in the cortex of normal MII oocytes (23, 38). The presence of similar SER-mitochondria clusters is documented by historical TEM studies (39–42). In light of these findings, it is tempting to speculate that the SER disc represents an extreme phenotype arising as the exaggeration of the physiological process taking place during oocyte maturation. The intimate bicomponent alliance of energy-supplying organelles and calcium-storing elements ensures effective calcium signaling required for oocyte activation (38, 43, 44). However, excessive SER aggregation reducing surface contact with mitochondria may diminish calcium availability during fertilization and thus compromise oocytés developmental competence.

Based on our ultrastructural data, refractile bodies and subtle SER aggregation could also occur in normal oocytes, whereas vacuolization appears to be an exclusively deviant feature. The variable number, size, and appearance of vacuoles support the hypothesis that these inclusions are products of rapid endocytic process and their content corresponds to perivitelline fluid (19). It is intuitive to presume that the larger portion of cytoplasm vacuoles occupy, the more severe the impact on oocytés fitness. In our settings, the large fluid-filled vacuoles posed a challenge for intracellular structure preservation, and some samples did not endure the fixation procedure. Similarly, the vacuolization may sensitize the oocyte to osmotic changes and hamper its cryosurvival. The simultaneous presence of giant vacuoles with large SER aggregates, and misshaped, most probably dysfunctional (45), mitochondria indicate that vacuolization can be a readily noticeable sign of major cellular disturbance. Horseshoe- and ring-shaped mitochondria have been previously detected in TEM micrographs of a SER-positive human oocyte and linked to oocyte deterioration (11).

Presented image data illustrated that the structural basis of CLCG and SER disc is anomalous aggregation of SER elements. This similarity, together with our clinical experience that both cytoplasmic texture irregularities typically occur in mature eggs and disappear upon sperm entry, suggests that the two phenotypes might have a common foundation. Biological nature, limited availability, and individual variability make functional experiments in human oocytes notoriously challenging. Here, we employed chemical inhibitors to address the role of the cytoskeleton in organelle arrangement in a representative number of human oocytes derived from young and healthy donors. Although selected drugs are commonly used in cytoskeleton research, their off-target effects can not be ruled out. Therefore, further experimental investigation is needed to validate our results. Nevertheless, the observed phenotype reproducibility in each treatment group and the number of controls support the relevance of presented data. Our finding that actin, not microtubules, plays a role in the homogenous distribution of organelles during human oocyte maturation contrasts with published evidence that microfilament depolymerization did not affect the motility of mitochondria in mouse and porcine oocytes (46, 47). On the other hand, the impact of both actin-targeting drugs on spindle-chromosome configuration is in line with the recognized role of spindle actin in human oocyte’s meiotic spindle assembly and chromosome alignment (48). The SER cisternae swelling observed in BFA-treated oocytes evidenced that the drug blocked vesicular trafficking from endoplasmic reticulum (ER) to the Golgi apparatus leading to the membrane recycling to ER. This phenomenon was particularly pronounced in GV oocytes and less apparent in maturing oocytes because the Golgi apparatus disassembles upon resumption of meiosis (23). Surprisingly, in our experiments, human oocytes exhibited insensitivity to mitotic poison nocodazole in a concentration capable of inhibiting spindle assembly in mouse oocytes (49). This observation suggests that human oocytes may be equipped with a mechanism for the efflux of specific foreign substances. The ability of human oocytes to actively eliminate xenobiotics was previously reported by Brayboy and colleagues (50), who speculated that protection against toxic substances might play a pivotal role in the survival of long-lived human oocytes exposed to environmental pollutants. Exploring oocyte chemoresistance might open new opportunities for developing therapeutical strategies to preserve the fertility of cancer patients.

Cytoplasmic abnormalities may indicate female gametes’ genetic, epigenetic, and metabolic defects. Large oocyte cytoplasm not only constitutes an intracellular signaling hub integrating external and internal cues, but also provides organelles, nutrients, and metabolites needed to support post-fertilization development. Alteration of cellular architecture might impact oocytés homeostasis, capacity to interact with sperm-delivered molecular factors, and sustain metabolism of early cleavage embryos. In transcriptionally-silent oocytes, the cytoplasm stores maternal mRNAs and ensures spatial and temporal control over the translation and degradation of transcripts synthesized during oogenesis (51). A recent study showed that mammalian oocytes, including humans, dock their maternal mRNA in mitochondria-associated membrane-less domains (52). Anomalous organelle distribution seen in aberrant oocytes is likely to compromise the availability of mRNA and, thus, the efficiency of protein synthesis and post-translation modifications. Since the cell mass of a newly formed zygote is almost exclusively maternally-inherited, a deficiency in ooplasmic properties would inevitably compromise the oocytés capacity to produce a viable embryo.

Objective oocyte rating would be especially useful in shared egg donor programs and medical freezing cycles. However, micrographs of abnormal morphotypes presented here, together with our earlier data from morphologically normal oocytes (23), indicate that routine stereoscopic examination is insufficient to assess egg quality. High magnification is needed to detect pathological features such as misshaped mitochondria and enlarged SER. The development of high-resolution tomographic methods now enables volumetric imaging of oocyte and ovarian tissue architecture (24, 53). Three-dimensional reconstruction of isolated and/or cumulus-enclosed dysmorphic oocytes would elucidate the spatial relationship of subcellular structures and advance our understanding of how the oocytes interact with surrounding cells. The unexplored character of the female gamete and its typical sphericity makes the commercial claims that oocyte ability to produce transferable embryos can be deduced from a single low-resolution snapshot questionable. The emergence of non-invasive label-free imaging techniques (54, 55) and omics approaches suitable to analyze individual oocytés microenvironment (56, 57) holds the potential to identify new biomarkers of egg quality for the benefit of IVF patients.

## CONCLUSIONS

This study enhances our understanding of the human oocytés internal organization and possible factors implicated in the evolution of certain morphological abnormalities. Presented micrographs collection extends rare documentation of the fine morphology and phenotypical variability of dysmorphic eggs. Furthermore, our inhibition experiments demonstrated that actin disruption leads to excessive organelle clustering and generates an ultrastructural pattern resembling naturally occurring aberrant phenotypes. Together, these data support the hypothesis that actin cytoskeleton dysfunction (along with other unknown factors) underlies oocyte dysmorphism encountered in the clinical IVF practice. We hope that this study’s data, along with those provided by other investigators, will contribute to forming the scientific groundwork for improved evaluation of egg morphology in ART.

## LIST OF ABBBREVIATIONS

ART: assisted reproduction technology
BFA: - Brefeldin A
CLCG: - centrally-located cytoplasmic granularity
COS: - controlled ovarian stimulation
CytD: - Cytochalasin D
DMSO: - dimethyl sulfoxide
GV: - germinal vesicle
ER: - endoplasmic reticulum
MII: - Metaphase II
mRNA: – messenger ribonucleic acid
ICSI: – intracytoplasmic sperm injection
IVF: - in vitro fertilization
NC: - Nocodazole PB - polar body
PBT: - phosphate-buffered saline supplemented with 0.1% Triton X-100
PLM: – polarized light microscopy
PVS: – perivitelline space PX – paclitaxel
SER: – smooth endoplasmic reticulum
TEM: – transmission electron microscopy
ZP: - zona pellucida

## DECLARATIONS

### Ethics approval

The study was undertaken under ethical approvals issued by the Ethics Committees of the collaborating academic institution (16/2016 and 1/2019) and clinical unit (1/2015). All donors provided written informed consent explaining the propose of the research and their engagement in the study.

### Consent for publication

All authors declare consent for publication.

### Availability of data and materials

Not applicable

### Authorś contribution

M.T.: Electron microscopy of dysmorphic oocytes, Inhibition experiments, Fluorescence microscopy, Data analysis, and interpretation, Manuscript and figure drafting; Z.T.: Sample processing, Electron microscopy of dysmorphic oocytes, Data analysis and interpretation; D.K.: Sample collection, Morphological assessment, Polarized Light Microscopy; P.O.: Hormonal stimulation, Recruitment of sample donors, Manuscript critical reading; S.K.: Informed consent collection and administration, Manuscript critical reading; Z.H.: Project design, Data analysis, and interpretation, Manuscript writing. All authors discussed the results and commented on the final manuscript.

## Acknowledgements

The authors wish to thank egg donors and clinical staff for providing research samples and administrating relevant documentation. We also acknowledge the core microscopic facility for technical support with the acquisition of fluorescent microscopy images presented in this paper.

## Authors’ information

Correspondence should be addressed to Zuzana Holubcová, Masaryk University Campus – building F01B1, Kamenice 3, 625 00 Brno, ORCID ID: 0000-0002-4658-6161

---

**Supplementary Figure 1:**
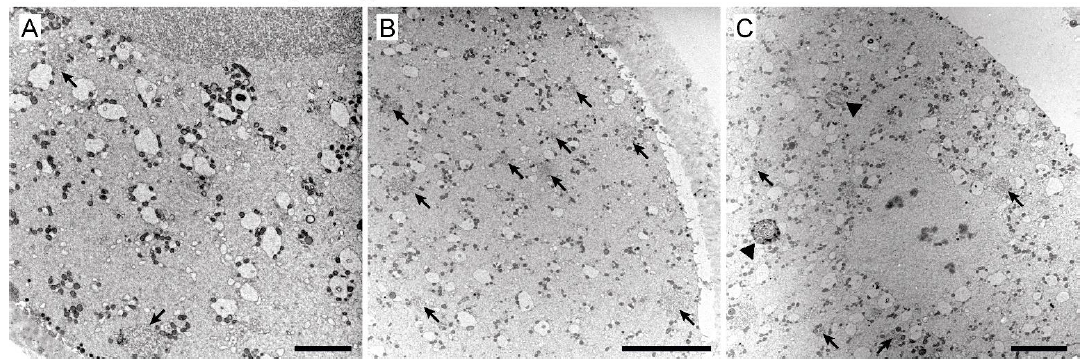
Representative TEM micrographs of SER disc oocytés cytoplasm. Mid-sized SER aggregates (A-C; arrows) and refractile bodies (C; arrowheads) are present in the ooplasm. Detail of meiotic spindle (C). The same sample as in Figure 3. Scale bar, 5 μm.

**Supplementary Figure 2:**
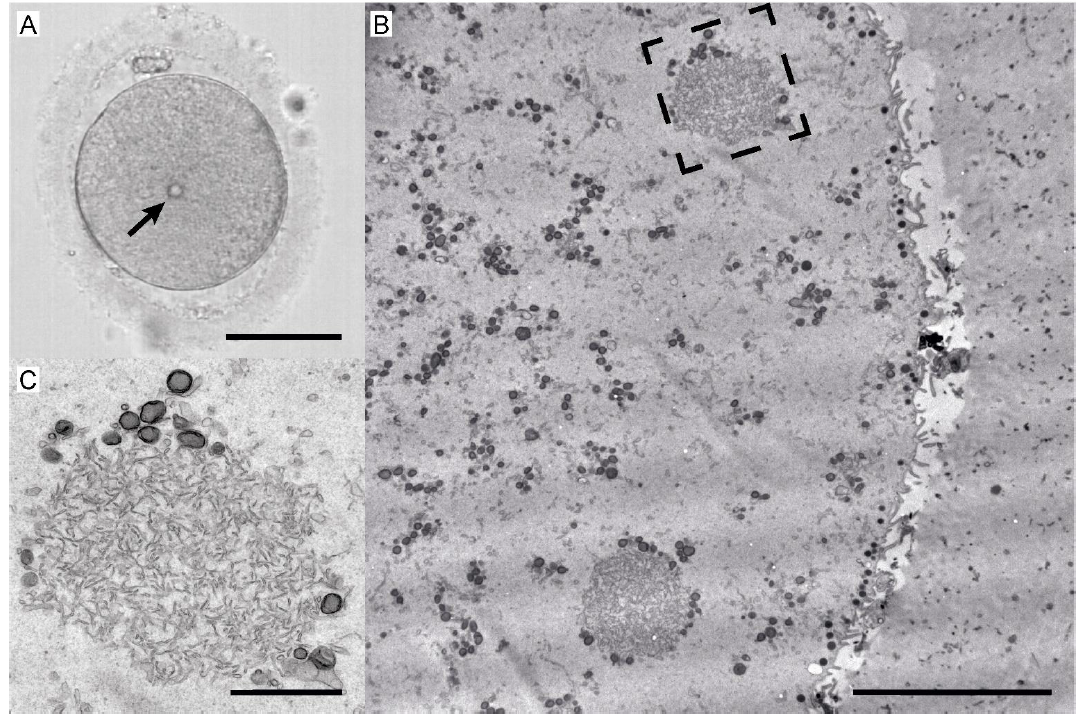
Example of the oocyte with mid-sized SER clusters undetectable by light microscopy. Live cell appearance in transmitted light (A) and TEM micrographs (B, C) of the oocyte exhibiting a prominent refractile body (arrow). Overview (B) and magnified detail (C; dashed rectangle in B) of enlarged SER clusters located in the cortical area are shown. Scale bar, 50 μm (A), 10 μm (B), 2 μm (C).

**Supplementary Figure 3:**
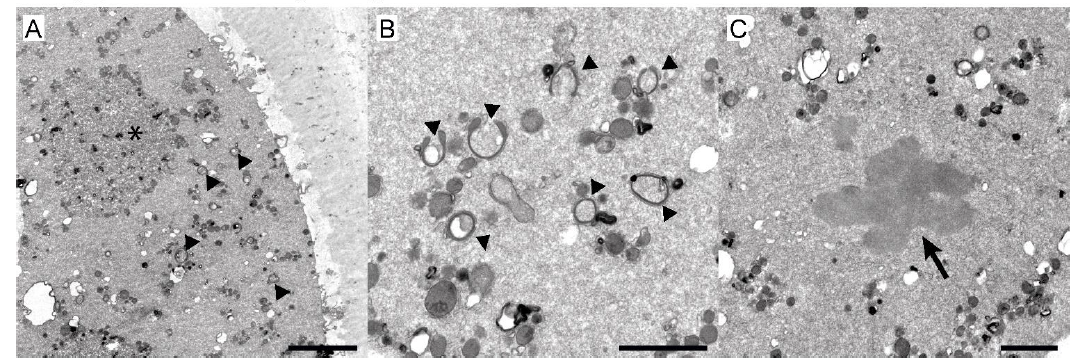
TEM micrographs of a vacuolized oocyte presenting mid-sized SER aggregate. Same sample as in Figure 4 A-C; asterisk indicates mid-sized SER aggregate (A), arrowheads indicate horseshoe- and ring-shaped mitochondria (A-C), and arrow indicates chromosome cluster (C). Numerous little vacuoles are visible in the cytoplasm (A-C). Scale bar, 5 μm (A), 2 μm (B, C).

**Supplementary Figure 4:**
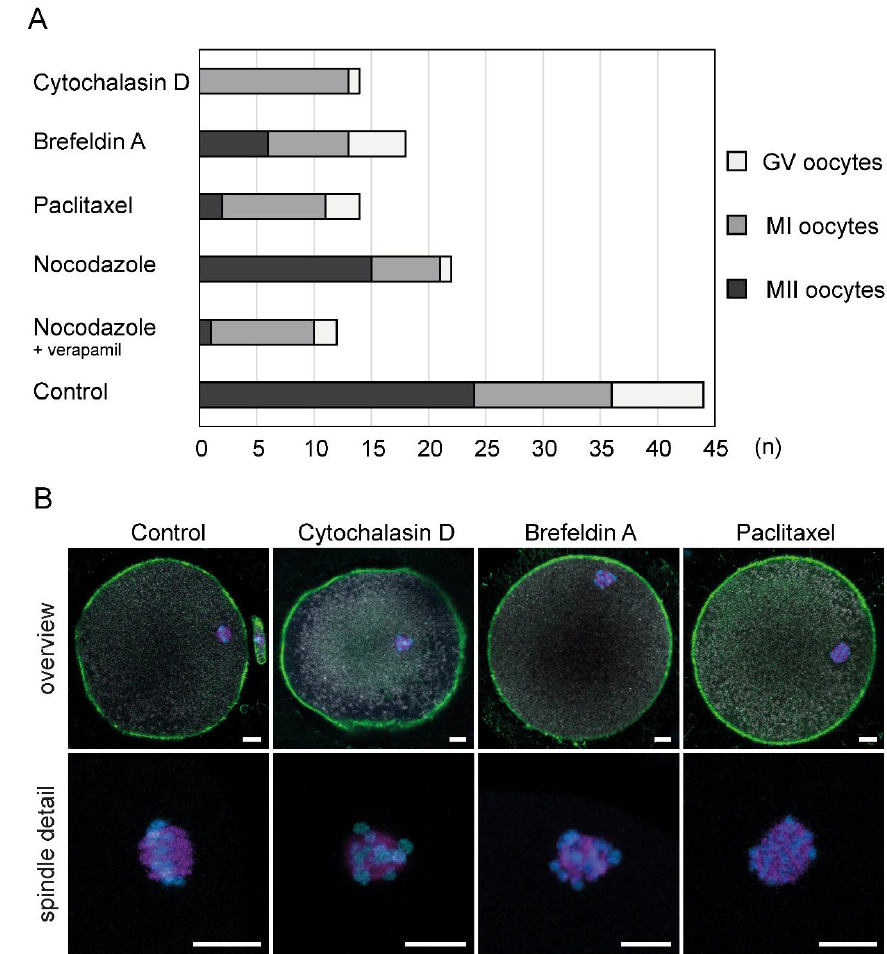
Maturation rate and morphology of drug-treated oocytes. Graph depicting in vitro maturation efficiency of drug-treated and control oocytes; n = number of oocytes (A). Fluorescence images of oocytes cultured overnight in control or experimental conditions (B). Overall cell morphology (top row) and meiotic spindle detail (bottom row) of a MII oocyte matured in presence of DMSO, and MI-arrested oocytes exposed to actin (cytochalasin D, brefeldin A) and microtubule (paclitaxel) inhibiting drugs. Actin microfilaments (green), microtubules (magenta), mitochondria (gray), and chromosomes (blue) are labeled. Scale bar, 10 μm.

**Supplementary Figure 5:**
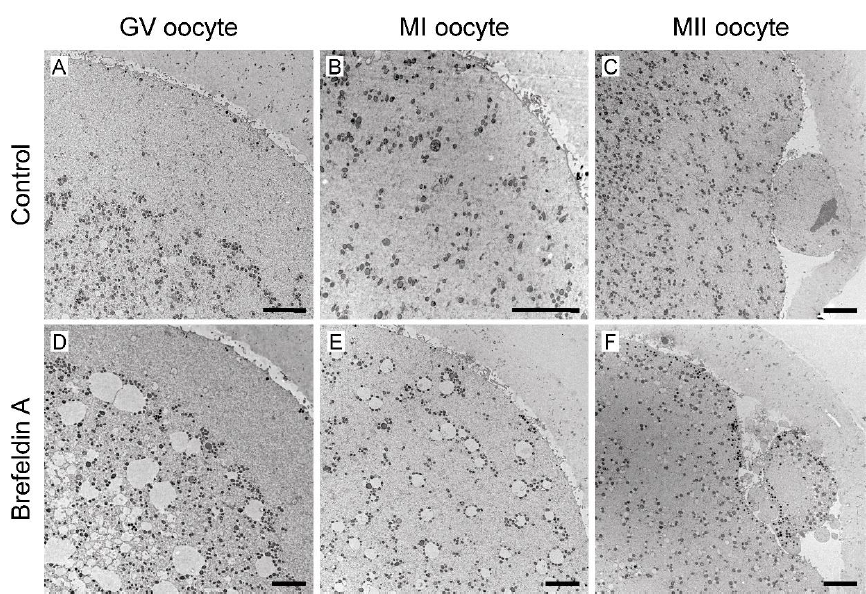
Effect of Brefeldin A exposure on intracellular morphology of human oocytes maturing in vitro. TEM micrographs of oocytes maturing in control conditions (A-C) and exposed to brefeldin A (D-F). The swelling of SER cisternae is prominent in brefeldin A-treated GV and MI (metaphase I) oocytes and diminishes in MII oocytes (D-F). Scale bar, 5 μm.

**Supplementary Figure 6:**
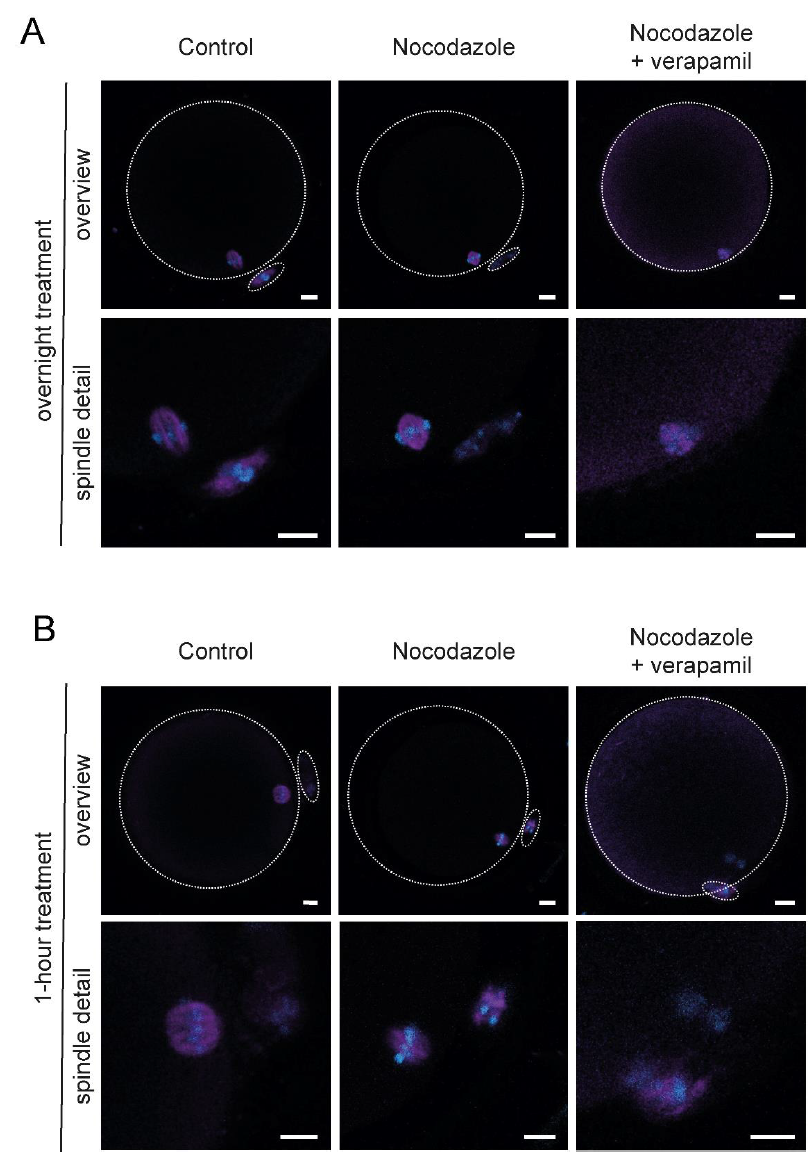
Short- and long-term treatment of human oocytes with nocodazole. Representative fluorescence images of oocytes exposed to DMSO (control) and nocodazole (+/− addition of verapamil) during overnight maturation from GV stage (A) and 1-hour treatment of MII arrested oocytes (B). Overviews (top row; cell outline is dashed) and spindle area details (bottom row) of oocytes with (immuno)labeled microtubules (magenta) and chromosomes (blue) are shown. Scale bar, 10 μm

**Supplementary Table.**
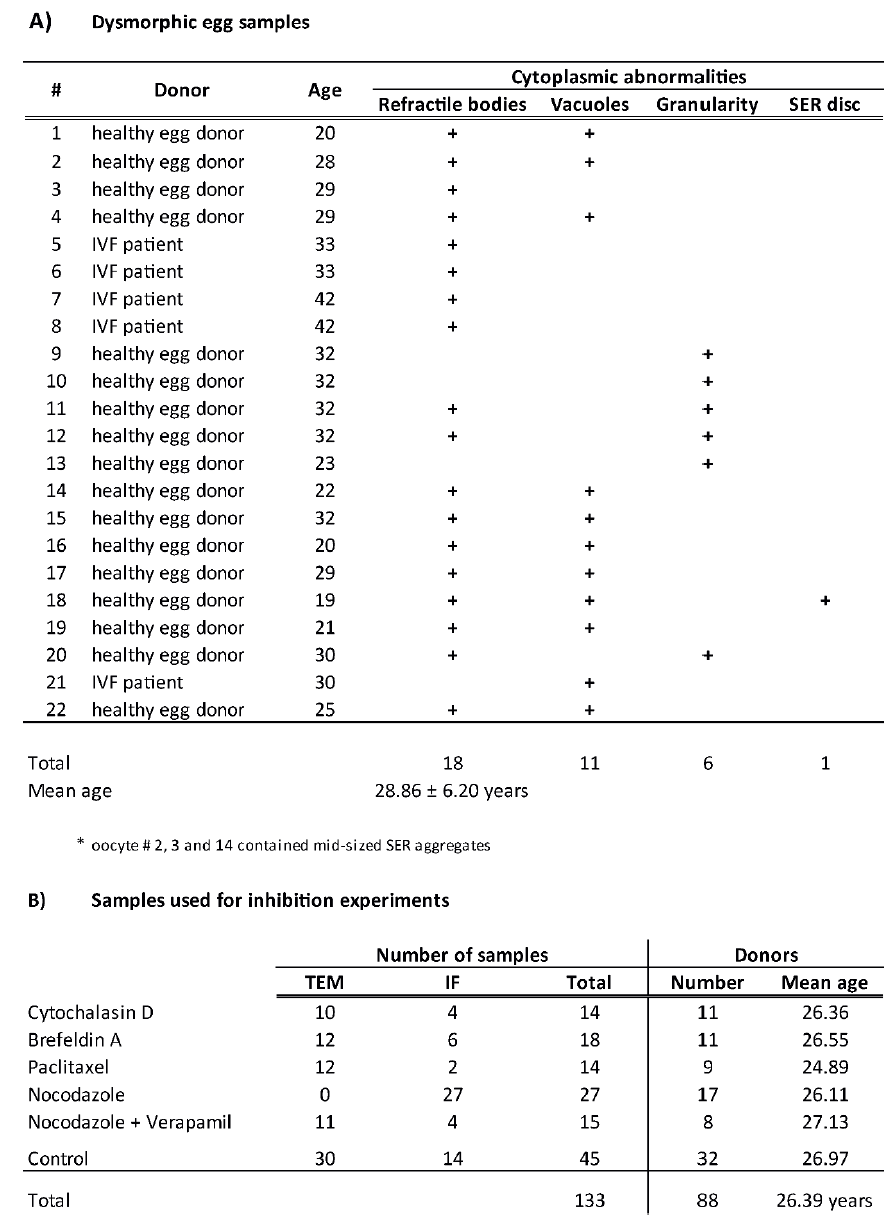
Overview of analyzed human oocyte samples.

A) Dysmorphic egg samples
| # | Donor | Age | Cytoplasmic abnormalities |  |  |  |
| --- | --- | --- | --- | --- | --- | --- |
|  |  |  | Refractile bodies | Vacuoles | Granularity | SER disc |
| 1 | healthy egg donor | 20 | + | + |  |  |
| 2 | healthy egg donor | 28 | + | + |  |  |
| 3 | healthy egg donor | 29 | + |  |  |  |
| 4 | healthy egg donor | 29 | + | + |  |  |
| 5 | IVF patient | 33 | + |  |  |  |
| 6 | IVF patient | 33 | + |  |  |  |
| 7 | IVF patient | 42 | + |  |  |  |
| 8 | IVF patient | 42 | + |  |  |  |
| 9 | healthy egg donor | 32 |  |  | + |  |
| 10 | healthy egg donor | 32 |  |  | + |  |
| 11 | healthy egg donor | 32 | + |  | + |  |
| 12 | healthy egg donor | 32 | + |  | + |  |
| 13 | healthy egg donor | 23 |  |  | + |  |
| 14 | healthy egg donor | 22 | + | + |  |  |
| 15 | healthy egg donor | 32 | + | + |  |  |
| 16 | healthy egg donor | 20 | + | + |  |  |
| 17 | healthy egg donor | 29 | + | + |  |  |
| 18 | healthy egg donor | 19 | + | + |  | + |
| 19 | healthy egg donor | 21 | + | + |  |  |
| 20 | healthy egg donor | 30 | + |  | + |  |
| 21 | IVF patient | 30 |  | + |  |  |
| 22 | healthy egg donor | 25 | + | + |  |  |
| Total |  |  | 18 | 11 | 6 | 1 |
| Mean age |  |  | 28.86 ± 6.20 years |  |  |  |

\* oocyte # 2, 3 and 14 contained mid-sized SER aggregates

B) Samples used for inhibition experiments
|  | Number of samples |  |  | Donors |  |
| --- | --- | --- | --- | --- | --- |
|  | TEM | IF | Total | Number | Mean age |
| Cytochalasin D | 10 | 4 | 14 | 11 | 26.36 |
| Brefeldin A | 12 | 6 | 18 | 11 | 26.55 |
| Paclitaxel | 12 | 2 | 14 | 9 | 24.89 |
| Nocodazole | 0 | 27 | 27 | 17 | 26.11 |
| Nocodazole + Verapamil | 11 | 4 | 15 | 8 | 27.13 |
| Control | 30 | 14 | 45 | 32 | 26.97 |
| Total |  |  | 133 | 88 | 26.39 years |

